# Longitudinal analysis of bacterial community dynamics in normal hospital ward following patient occupancy

**DOI:** 10.1101/2025.09.18.677220

**Authors:** Liyun An, Xiaonan Liu, Xiao Li, Yixuan Chu, Qiang Li, Xiaoqiang Sun, Junsheng Chu, Yong Nie

**Author notes:** Corresponding authors: *E-mail addresses:* (J. Chu), (Y. Nie).

## Abstract

The hospital microbiome significantly influences patient recovery and clinical outcomes. However, the dynamics of microbial colonization and transmission following initial patient occupancy remain poorly understood. Here, we employed 16S rRNA gene amplicon sequencing to investigate bacterial community dynamics on surfaces within neurosurgery ward and patients as a new hospital became operational. Our results showed that bacterial colonization in hospital wards follows distinct site-specific patterns, after hospital opening, alpha diversity was significantly increased on floors and drawer handles but decreased on bedrails and faucet handles compared to preopening. Beta diversity analysis showed that surfaces frequently contacted by patients exhibited the greatest compositional turnover, such as bedrails, drawer handles, and faucet handles, bacterial communities in after-opening were more homogeneous across sites than preopening, indicating potential bacterial transmission. Moreover, we found following patient admission, patient hand-derived microbiomes exert a significant influence on the bacterial communities in hospital wards, with a particularly pronounced impact on bedrails. Additionally, the potential disease risk of bedrails in post-opening was significantly higher than preopening. Taken together, these findings underscore the critical role of human contact in shaping hospital microbiomes and highlight the importance of targeted infection control strategies to mitigate pathogen transmission.

## Introduction

Interactions between environmental microbiomes and humans play a critical role in human health and disease^1^. As urbanization progresses, individuals spend more time in shared public building environments with strangers^2,3^, such as offices, schools, and hospitals. The shared occupancy not only contributes to the complexity of indoor microbial communities but also increases the risk of exposure to a wide range of microorganisms, including potential pathogens^4^. Consequently, in recent decades, a growing number of studies have explored the interactions between human-associated microbiomes and built environments, including museums^5^, kindergarten^6^, shopping malls^7^, public transit systems^8,9^, and hospitals^10–13^.

The hospital ward is a special public indoor environment in which microorganisms continuously interact with patients, thereby exerting a direct influence on patient recovery and clinical outcomes^14^. Although clinical environments are subject to stringent hygiene guidelines^15,16^, freshly cleaned, sanitized, or sterilized surfaces possess only a transient aseptic state^17^. Therefore, hospital-acquired infections (HAIs) in medical care remain a major threat to patient life and are now recognized as one of the most serious public health challenges of the 21^st^ century, such as leading to increased morbidity and mortality, prolonged hospital stays, and greater financial burdens on both healthcare systems and families^18^. In the United States, it is estimated that one in every 31 hospitalized patients suffers from at least one HAI every day^19^.

Preventing HAIs requires a comprehensive understanding of hospital microbial diversity, as well as the identification of sources and transmission pathways of infectious agents^20,21^. Emerging evidence indicates that environmental contamination and opportunistic pathogens in clinical settings substantially contribute to HAI development^2,18,22,23^. Certain pathogens are particularly well-adapted to the hospital environment and can colonize surfaces that frequently come into close contact with patients or medical equipment^23–25^, thereby substantially increasing the risk of pathogen transmission. Previous studies have implicated that hospital ward environment is an important reservoir for colonizing bacteria, and high-touch surfaces such as bedrails, floors, and handles are especially susceptible to microbial contamination^26^. Therefore, understanding the hospital microbiome is essential not only for maintaining low levels HAI incidence, but also for enhancing the overall quality of healthcare delivery.

A larger-scale analysis of hospital microbial communities is critical for understanding how microorganisms spread within hospital environments and to what extent the skin microbiomes of patients influence and are influenced by their surroundings. Apart from a limited number of studies confined to intensive care units (ICUs)^10,27–33^, to date, there are relatively few detailed longitudinal, culture-independent studies on the overall microbiome composition in normal hospital. Additionally, newly constructed building provides unique opportunities to investigate the temporal dynamics of bacterial communities. For example, Lax et al. conducted a landmark study that characterized the microbial ecology, colonization patterns, and succession within a newly built hospital, and identified distinct ecological signatures of bacterial exchange among the environment, patients, and healthcare workers^26^. In detail, these pioneer studies found that surfaces such as bedrails and hot water faucets were among the most active zones of microbial transfer, while floors demonstrated comparatively static microbial communities^26^. Another longitudinal study investigated the dynamics of dust microbiome in a newly opened kindergarten and found significant temporal shifts in microbial community structure, particularly in the biomass of human-associated bacteria taxa^6^. Recent surveillance data have reported high incidence of HAIs in normal hospital wards, including neurology stations^11,26^, underscoring the need to expand microbiome research beyond ICUs. However, the processes of colonization and transmission following hospital opening and the admission of the first patients remain poorly understood. A deeper understanding of the microbial colonization dynamics and transmission pathways within hospital environments is essential for developing innovative strategies to mitigate the spread of nosocomial pathogens, minimize patient co-infection risk, and ultimately optimize healthcare-system efficiency. Therefore, further research into the colonization and movement of bacteria in hospital environments is not only warranted but necessary for improving patient outcomes and optimizing clinical hygiene protocols.

In this study, we investigated the microbial community profiles within the neurosurgery ward environment of the newly constructed Beijing Tiantan Hospital in China. Samples were collected from a variety of surfaces, including floors, bedrails, drawer handles, faucet handles, and from patients. This study aimed to: (1) describe the alterations in colonizing bacterial communities after hospital opening, (2) clarify bacterial transmission patterns within hospital wards, and (3) explore changes in potential disease risks and the susceptible human organs after hospital opening. The results of this work reveal the characteristics of pre-opening microbiome and its compositional shifts following patient occupancy, offering insights into bacterial transmission dynamics within clinical environments, and providing valuable guidance for healthcare management and the prevention of co-infections.

## Methods

### 1. Sample collection and DNA extraction

To investigate how microorganisms colonize and spread in the hospital environments, we conducted a five-month sampling from the opening of the hospital to its normal operation. In total, 270 hospital environmental samples and 72 patient samples were collected from the neurosurgery ward in newly constructed Beijing Tiantan Hospital, China, including floors, bedrails, drawer handles, faucet handles, patient hands, patient noses, and patient axillae (Table S1). These surfaces are frequently touched by patients and healthcare workers, making them focal points for targeted cleaning and disinfection to minimize the risk of pathogen transmission. All samples were collected using a nylon flocked swab (COPAN, 5U018S.CN) premoistened in sterile buffer (0.15 M NaCl, 0.1% Tween 20). For floor sampling, an area of approximately (10 cm × 20 cm) was swabbed for 30 s. For bedrail sampling, the top and side surface at midpoint of bedrails were swabbed for 30 s. For drawer handle and faucet handle sampling, the top, middle, and bottom of each handle were swabbed for 30 s. After sampling, to avoid hand contamination, the swab tips were cut off using sterile scissors, placed in the sealed transport tubes, immediately transported to the laboratory on ice and stored at −80°C until DNA extraction. Total DNA was extracted using the DNeasy^®^ PowerSoil^®^ kit (QIAGEN, Germany). Firstly, the swab tips were transferred into a PowerBead Tube provided in the kit, subsequently, the tube was vortexed for 30 s, and the tip was then discarded. DNA was then extracted according to the manufacturer’s instructions.

### 2. 16S rRNA gene sequencing and processing

The V4 hypervariable region of the 16S rRNA gene was amplified by polymerase chain reaction (PCR) using the 515F (5′-GTGYCAGCMGCCGCGGTAA-3′) and 806R (5′-GGACTACNVGGGTWTCTAAT-3′) primers. Each primer contained an Illumina-specific adaptor and a unique barcode, enabling multiple samples to be pooled within a single sequencing run. PCR amplification was conducted in a 50 μL reaction mixture containing 25 μL of 2 × Premix Taq, 1 μL of forward primer (10 μM), 1 μL of reverse primer (10 μM), 50 ng of template DNA, and nuclease-free water. The PCR cycling conditions were as follows: an initial denaturation at 94°C for 5 min; followed by 30 cycles of denaturation at 94°C for 30 s, annealing at 52°C for 30 s, and extension at 72°C for 30 s; and a final extension at 72°C for 10 min to ensure complete amplification. Sequencing was then performed on the Illumina HiSeq platform at Guangdong Magigene Biotechnology Co., Ltd. China, as previously described^34^.

The raw sequencing reads were processed using the QIIME 2 pipeline (v2019.7)^35^. Barcodes and primers were trimmed with the Cutadapt plugin^36^. Clean reads were then denoised and clustered into amplicon sequence variants (ASVs) using the DADA2 pipeline^37^. Taxonomy classification of ASVs was conducted with the SILVA database (release 128)^38^. Sequences identified as bacteria origin were retained. Additionally, singleton ASV was discarded. All samples were rarefied to a minimum number of sequences from each sample (46,352 reads). Bray-Curtis distance and alpha diversity metrics, including Shannon, richness, and phylogenetic diversity (Faith’s PD), were calculated using the diversity plugin.

The raw reads were deposited into the NCBI Sequence Read Archive (SRA) database (Accession Number: PRJCA046001).

### 3. Network analysis, source tracking analysis, and microbial index of pathogenic bacteria

To minimize artificial associations among low-abundance bacteria and simplify network complexity, only genera with an average relative abundance greater than 0.001% across all samples were retained for network construction^39^. Pairwise Spearman’s correlations between genera were computed, and relationships were considered valid when correlation coefficient was greater than | 0.7 | and adjusted P-value (Benjamini and Hochberg) was less than 0.01. Within the network, each node represents one genus, and each edge indicates a statistically significant association between nodes. Networks visualization was performed using the Gephi platform^40^. Additionally, node-level topological features of the networks were calculated to estimate the importance of each node, including degree, betweenness, closeness, and eigenvector centrality.

Bacterial transmission routes within hospital wards were determined using SourceTracker software package^41^, which employs a Bayesian model in combination with Gibbs sampling to quantify the proportion contribution of taxa from multiple source environments contributes to a sink environment. The transmission of bacteria across different media within hospital setting is complex and challenging to characterize. Therefore, in order to highlight critical transmission chains, we constructed a "Source to Sink" framework that encompassed all possible transmission directions in hospital wards. The contribution values of source tracking task served as quantitative indices to present the microbial transfer potential from a single “source” site to a “query” site. Based on these contribution values, we mapped the microbial transmission network within hospital ward by Cytoscape v3.5.1^42^ (http://cytoscape.org).

Microbial Index of Pathogenic bacteria (MIP) measures the potential pathogenicity of the microbial community. We used Parallel-MeTA3 (https://github.com/qdu-bioinfo/mip) to calculate MIP value based on the cumulative relative abundance of all opportunistic pathogenic bacteria in bacterial community. MIP value ranged from 0 to 1.

### 4. Statistical analysis

All statistical analyses were conducted in R (version 4.0.0)^43^. Sample size was not predetermined through statistical methods, and all collected data were included in the analyses. For beta diversity, Principal Coordinates Analysis (PCoA) was performed based on Bray-Curtis distance derived from ASVs abundance tables using the “vegdist” function in the vegan package^44^. To further assess compositional differences in bacterial communities across sites, we conducted a similarity analysis (ANOSIM) using the “anosim” function in the vegan package. All difference analysis between groups were calculated using Wilcoxon rank-sum test. Visualization was conducted using the ggplot2 package^45^, where stacked bar charts were constructed with “geom_col”, boxplots with “geom_boxplot”, and scatter plots with “geom_point” function.

## Results

### 1. Overall bacterial community overview of the hospital wards

To gain a comprehensive understanding of the bacterial community in hospital wards, we investigated both community composition and diversity indices. Phylum-level taxonomic assignment showed that Proteobacteria, Firmicutes, Actinobacteria, and Bacteroidetes were highly abundant across all four types of hospital ward samples (Fig. 1A), which is similar to previous research in hospital surfaces^26^. The average relative abundance sums of these four phyla were 92.44%. At the genus level, *Vibrio*, *Acinetobacter*, *Staphylococcus*, *Corynebacterium*, and *Pseudoalteromonas* were identified as dominant across all sample types. Additionally, *Enhydrobacter* were found to be particularly abundant in faucet handles (Fig. 1B).

**Fig. 1.**
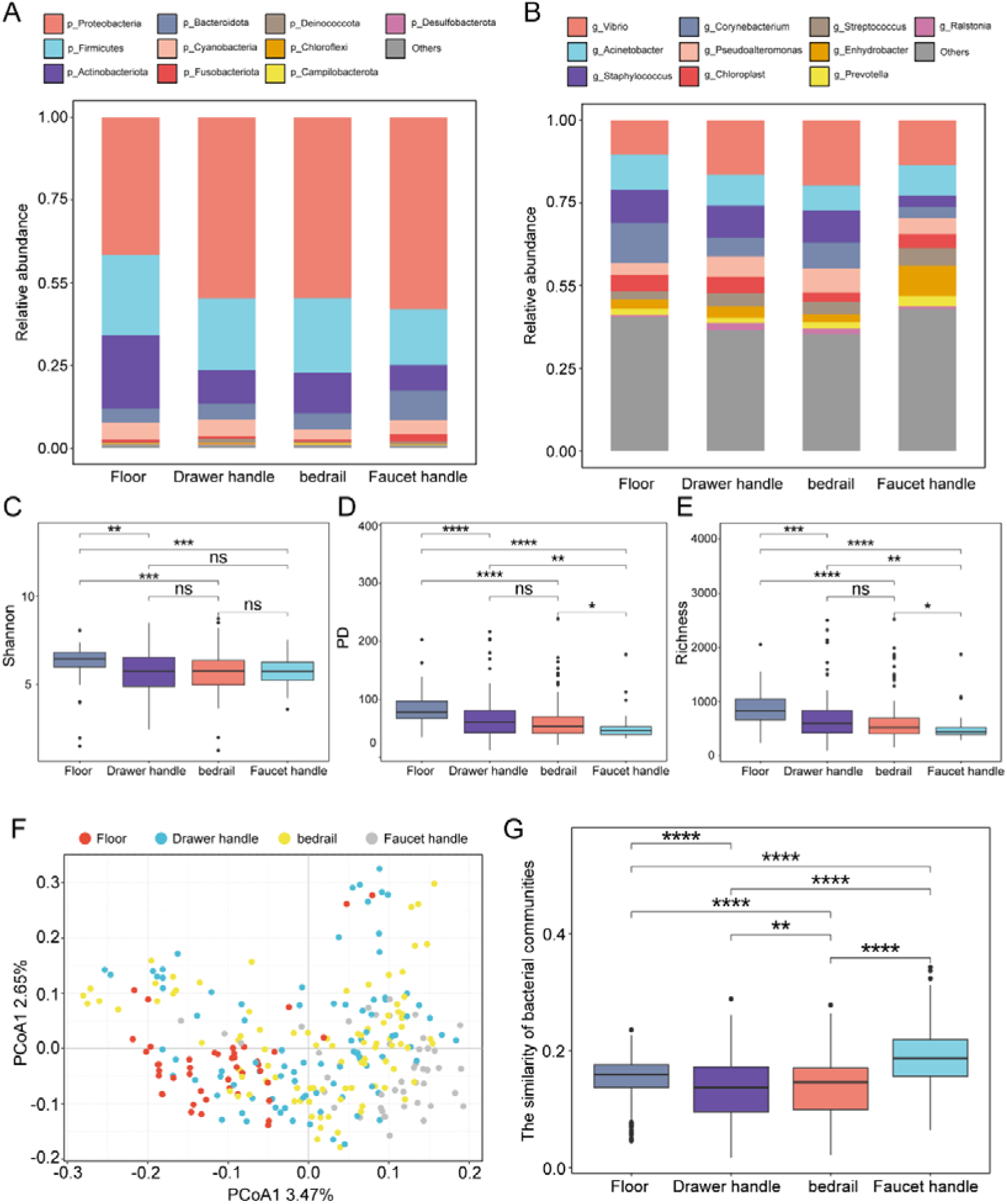
Bacterial composition and diversity metrics of hospital ward samples. (A, B) The composition of bacterial community at phylum (A) and genus (B) level. (C, D, E) The alpha diversity indices of environmental samples, including Shannon (C), Faith’s PD (D), and richness (E). Alpha diversity of bacterial communities in floors was higher than other samples. (F) Principal coordinates analysis (PCoA) of bacterial communities using Bray-Curtis distance. (G) The similarity of bacterial communities (1-Bray-Curtis distance) in various hospital ward samples.

Furthermore, we compared the diversity of the bacterial communities across various hospital ward samples. The results showed that floor samples exhibited the highest bacterial diversity (Fig. 1C, D, E), likely reflecting their frequent exposure to diverse external microbial inputs due to direct or indirect contact with the outdoor environment. In addition, PCoA based on Bray-Curtis distance showed that floor, bedrail, drawer handle, and faucet handle samples segregated into distinct clusters within the ordination space, ANOSIM test further confirmed significant differences among these clusters at the ASV level (Fig. 1F, ANOSIM, R = 0.06, *P* = 0.004). In addition, to further explore intra-group bacterial community similarity, we calculated pairwise bacterial community similarities at the ASV level, and found that faucet handle samples exhibited the highest within-group similarity compared to other sample types (Wilcox test, *P* < 2.2e-16; Fig. 1G). These results suggested that colonization process of bacterial communities in hospital wards follows a site-specific pattern, potentially shaped by differential exposure, human contact, and surface characteristics.

### 2. Bacterial colonization dynamics of the hospital wards

We next explored the bacterial colonization dynamics within hospital wards. When observing the average relative abundances at each date during the sampling, we observed rapid taxonomic successions in microbial community composition over time. In detail, at the phylum level, the bacterial communities across various hospital ward samples were primarily dominated by Proteobacteria, Firmicutes, and Actinobacteria both in the pre-opening and after-opening periods. The relative abundance of Proteobacteria decreased on both floors and drawer handles after the hospital opened, while the relative abundances of Firmicutes and Actinobacteria increased. In contrast, the relative abundance of these dominant phyla remained relatively stable on the bedrails and faucet handles (Fig. 2A, B). At the genus level, the bacterial communities in different sites of hospital wards were predominantly dominated by *Vibrio*, *Acinetobacter*, *Staphylococcus*, *Corynebacterium*, and *Pseudoalteromonas* both in the preopening and after-opening periods. As soon as the hospital became operational, the floors, drawer handles, and bedrails displayed an increased relative abundance of human skin-associated bacterial genera, including *Corynebacterium*, *Staphylococcus*, and *Streptococcus* (Supplementary Fig. 1A, B).

**Fig. 2.**
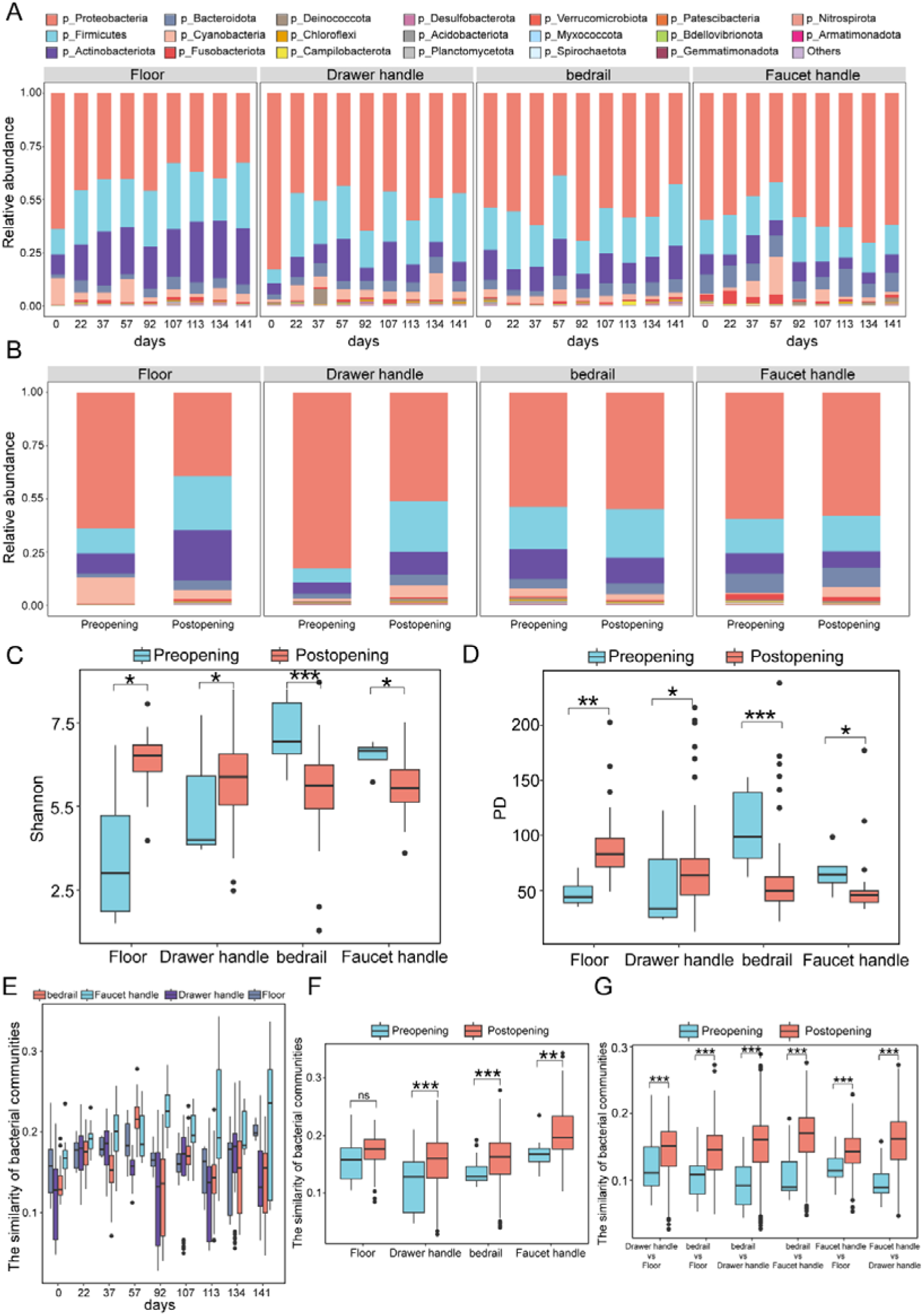
Change in bacterial community composition and structure after hospital opening. (A, B) Changes in the relative abundance of top10 phylum after hospital opening. (C, D) Box plot of changes in the Shannon diversity (C) and Faith’s PD (D)index of samples after hospital opening. (E) Change in the bacterial community similarity after hospital opening. (F, G) The comparison of bacterial community similarity between same (F) or different (G) sites in the pre-opening and post-opening.

Alpha diversity analysis showed that both Shannon diversity and PD index were significantly different between preopening and after-opening periods (Fig. 2C, D). Specially, we found that the alpha diversity of floors and drawer handles increased after hospital opened (Supplementary Fig. 2A, B, E, F), floors and drawer handles had a significantly lower alpha diversity in preopening compared to after-opening (Fig. 2C, D). Conversely, bedrails and faucet handles exhibited a decline in alpha diversity after hospital opened (Supplementary Fig. 2C, D, G, H), the alpha diversity was significantly higher in preopening than after-opening (Fig. 2C, D). The beta-diversity analysis at each site showed a rapid microbial colonization process. After hospital opened, the bacterial communities on faucet handles exhibited the most pronounced changes compared to other sites, with intra-group bacterial community similarity initially increased and then stabilized (Fig. 2E). Additionally, the intra-group bacterial community similarity of the floor, drawer handle, and bedrail samples showed an initial increase followed by a subsequent decline (Fig. 2E). Comparative analysis of beta diversity between the preopening and post-opening revealed that the intra-group bacterial community similarity in preopening was significantly lower than post-opening, with the exception of the floors (Fig. 2F). Moreover, inter-group bacterial community similarity in post-opening was significantly higher than preopening (Fig. 2G). The greatest differences in inter-group bacterial community similarity were observed between faucet handles and drawer handles, faucet handles and bedrails, and bedrails and drawer handles (Fig. 2G), likely due to increased contact frequency following the opening. Collectively, these results suggested that bedrails, drawer handles, and faucet handles represent the most dynamic habitats for microbial transfer, whereas floors serve as the least dynamic.

### 3. Hospital ward bacterial cohlonization dynamics shaped by patient occupancy

To highlight the potential influence of human activity on the bacterial composition of hospital wards, we investigated microbial exchange and interaction between surrounding environmental surfaces and patient-associated sites. In total, we analyzed 72 patient samples collected from three body sites over a five-month period after hospital opening, including nose (n = 24), hand (n = 24), and axilla (n = 24). Our analysis found that microbial communities across these different body sites of the patient were both diverse and abundant. At the phylum level, the hand, axilla, and nose samples were both dominated by Firmicutes, Proteobacteria, and Actinobacteria (Fig. 3A). In the axilla and nose samples, Firmicutes was the most abundant phylum, while in the hand samples, Proteobacteria was the most dominant phylum, with Firmicutes being the second most abundant (Fig. 3A). At the genus level, the hand, nose, and axilla samples were both dominated by *Staphylococcus*, *Corynebacterium*, and *Vibrio* (Fig. 3B). The relative abundance of *Staphylococcus* and *Corynebacterium* in axilla samples were both higher than in either hand or nose samples (Fig. 3B). Hand samples had the greatest alpha diversity, followed by nose samples, while axilla samples had the lowest (Fig. 3C, D).

**Fig. 3.**
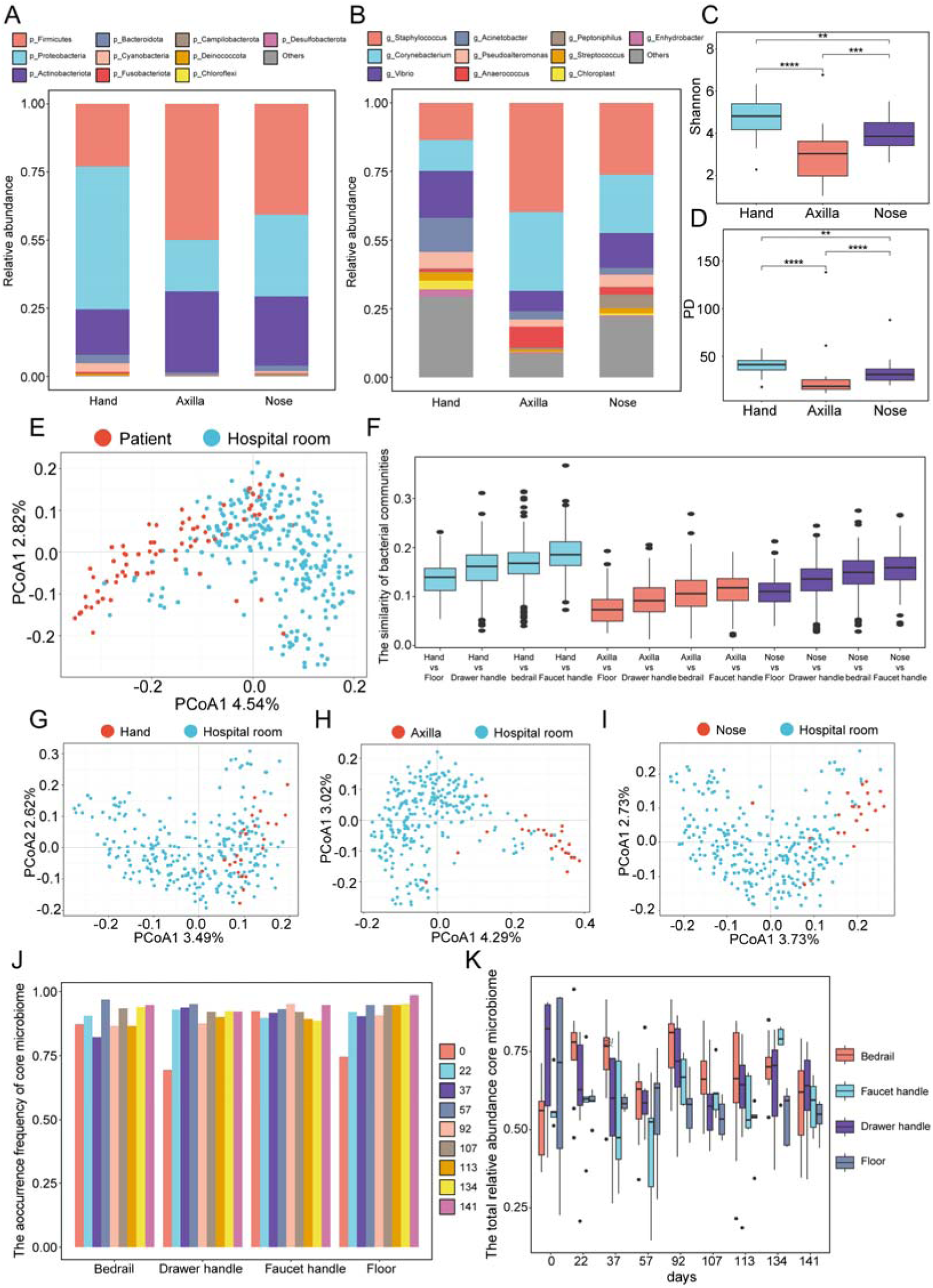
Bacterial composition and diversity metrics of patient samples. (A, B) The composition of bacterial community at phylum (A) and genus (B) level. (C, D) Shannon (C) and Faith’s PD (D) index of patient samples. (E, G, H, I) Principal coordinates analysis (PCoA) of bacterial communities using Bray-Curtis distance. (F) The similarity of bacterial communities (1−Bray-Curtis distance) between hospital wards and patients. (J, K) Change in the occurrence frequency (J) and total relative abundance (K) of core microbiome from patient hands in hospital wards after hospital opening.

Joint analysis of the four hospital ward sample types and the three patient body site samples in the PCoA space demonstrated a significant segregation between the patient-associated and the environmental cluster (Fig. 3E, ANOSIM, R = 0.16, *P* = 0.001). This separation indicated that the bacterial communities on different patient body sites were markedly distinct from those on various environmental surfaces within hospital wards. To quantify the differences in bacterial community composition across sampling sites, we conducted pairwise comparisons using Bray-Curtis distance and plotted as boxplot (Fig. 3F). The results showed that the bacterial communities on patient hands exhibited the highest similarity to those found on hospital wards, followed by noses, while axilla samples showed the greatest dissimilarity from environmental sites (Fig. 3F). Moreover, through PCoA analysis, we further elucidated that the bacterial communities on patient hands were most closely related to those on hospital wards (Fig. 3G, H, I). These results suggested that patient hands were the primary vector for microbial transfer between patients and hospital wards.

To further assess bacterial transmission and colonization of patient hands in hospital wards after hospital opened, we analyzed the distribution pattern of the core microbiome from patient hands across various sampling sites. The result showed that the occurrence frequency of core microbiome from patient hands in drawer handles and floors were obviously increased after hospital opened (Fig. 3J), indicating significant microbial transmission between patient hands, drawer handles, and floors. The occurrence frequency of core microbiome from patient hands in bedrails and faucet handles consistently remained high both before and after the hospital opening. Furthermore, we found the total relative abundance of core microbiome from the patient hands in drawer handles and floors were decrease after hospital opened, while a marked increase in bedrails (Fig. 3K). These results suggested that while microbes from patient hands spread widely in drawer handles and floors after the hospital opened, their colonization ability was weak. In contrast, bacterial colonization in bedrails by patient hands was most extensive. In summary, following patient admission, the patient hand-derived microbiomes exert a significant influence on the bacterial communities in hospital wards, with a particularly pronounced impact on bedrails.

### 4. Potential patterns of microbial transmission and Transfer-Easy taxa in hospital wards

Although the potential risk of microbial transmission within hospital wards has been well recognized, the bacterial transmission patterns remain unclear. In this study, we investigated microbial transmission by performing microbial source tracking. As shown in Fig. 4A, in the hospital wards, bidirectional bacterial transmission was observed between patients and hospital wards, patient hands, bedrails, and drawer handles served as major hubs for bacterial transmission. The majority of bacteria were transmitted from patient hands to noses, the remaining bacteria were transferred to bedrails and subsequently from bedrails to various other sites, such as floors, drawer handles, and faucet handles. Taken together, these findings elucidated the primary bacterial transmission routes across multiple sites, offering valuable guidance for hospital ward cleaning and management.

**Fig. 4.**
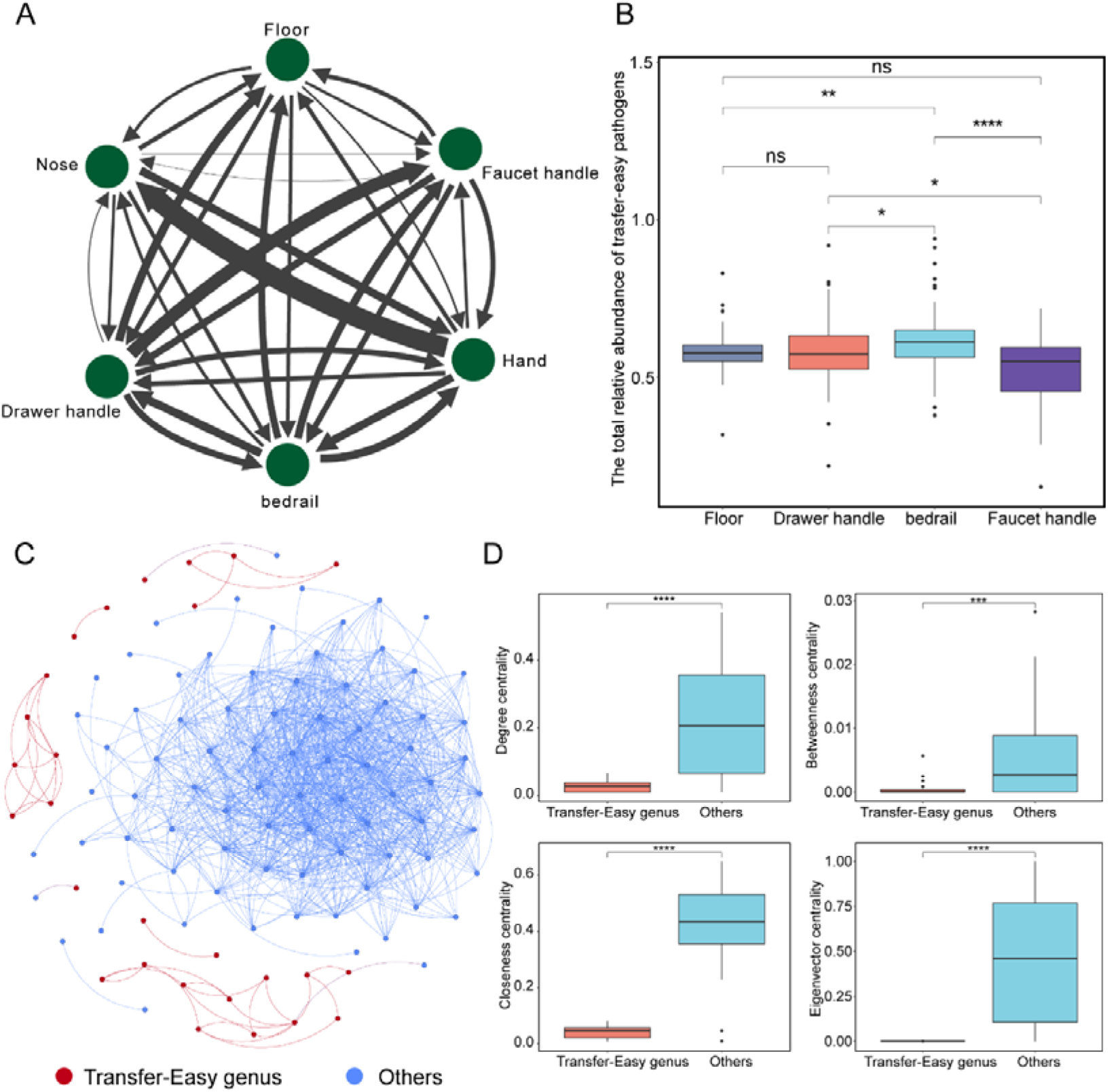
Microbial source tracking network and co-occurrence network of bacteria in hospital wards. (A) Network shows the source tracking contribution between patients and hospital wards. The contribution values from source sites to query sites were represented by the width of the edges. (B) The relative abundance of transfer easy pathogenic genus in hospital wards. (C) The co-occurrence network of bacterial communities in hospital wards. Nodes are colored based on transfer-easy genera and other genera. Each edges represent significant correlations (|ρ| > 0.7, *P* < 0.01). (D) Comparative analysis of node-level topological features between transfer-easy genera and other genera based on Wilcoxon rank sum tests, including degree, betweenness, closeness, and eigenvector centrality (\**P* < 0.05; \*\**P* < 0.01; \*\*\**P* < 0.001.

Various bacteria in hospital wards exhibit different transmission capabilities. In this study, we focused on bacteria that were prone to transmission within hospital wards, especially those disseminated to patients. We defined “Transfer-Easy taxa (TE taxa)” as taxa that detected in ≥ 80% of samples. In the hospital wards, a total of 71 genera were identified as TE taxa, among which 33 genera contain species previously reported to be pathogenic. The relative abundance of these pathogenic genera in bedrails was significantly higher compared to other sites within hospital wards (Fig. 4B). To further explore the ecological roles of TE taxa within the bacterial communities, we constructed co-occurrence network inferred from pairwise correlations, identifying the position of TE taxa within the network, and evaluated their relative importance through comparative analyses of node-level topological features between TE taxa nodes and other nodes. The result showed that the interactions among TE taxa were frequent, and only a minority of TE taxa had connections to other nodes (Fig. 4C). Additionally, we found node level topological features were both significantly lower (*P* < 0.001) for TE taxa than other nodes, including degree, betweenness, closeness, and eigenvector centrality (Fig. 4D). These results suggested that TE taxa possessed less connections with other nodes in hospital wards, implying that the transmission of TE taxa in hospital wards may not rely on interactions with other bacteria. Instead, it likely occurs through independent or direct routes, such as physical contact or environmental transmission. Therefore, efforts to control the spread of TE taxa should focus on strategies such as physical isolation and environmental disinfection, rather than on regulating microbial communities.

### 5. Potential disease risks and susceptible human organs in hospital wards

Moreover, to evaluate the potential disease risks and identify vulnerable human organs in hospital wards, we calculated the microbial index of pathogenic bacteria (MIP), which detects pathogenic bacteria in the microbiome and provides insights into disease risks and organ susceptibility. As shown in Fig. 5A, the overall MIP value was highest for floors, followed by bedrails, while faucet handles exhibited the lowest MIP values. Secondly, by calculating the relative contribution of individual species to the overall MIP value, we identified the top five pathogens that pose potential disease risks across various sites in hospital wards, including *Staphylococcus epidermidis*, *Acinetobacter baumannii*, *Klebsiella pneumoniae*, *Prevotella melaninogenica*, and *Streptococcus agalactiae* (Supplementary Fig. 3). Apart from *Streptococcus agalactiae*, the remaining four pathogens were classified as easily transferred pathogens. In addition, the average relative abundance of *Acinetobacter baumannii* and *Staphylococcus epidermidis* was notably high, at approximately 5%. These results indicated that hospital wards present a serious risk of transmission and colonization of pathogens. Moreover, we found pathogens cause diseases in human organs, such as oral and five sensory organs (OFSO), skin (SK), circulatory system (CS), urogenital system (US), digestive system (DS), respiratory system (RS), and others (e.g., motor system, nervous system, and endocrine system, OT) (Fig. 5B). Among them, MIP values related to CS diseases were highest in different sites of hospital wards (Fig. 5B), further analysis revealed that potential disease risks related to the CS mainly include septicemia, endocarditis, and blood stream infection (Fig. 5C). Additionally, given the samples analyzed in this study were collected from neurosurgery ward, diseases related to nervous system were also analysis and the result showed that potential disease risks related to the nervous system mainly include meningitis (Fig. 5D).

**Fig. 5.**
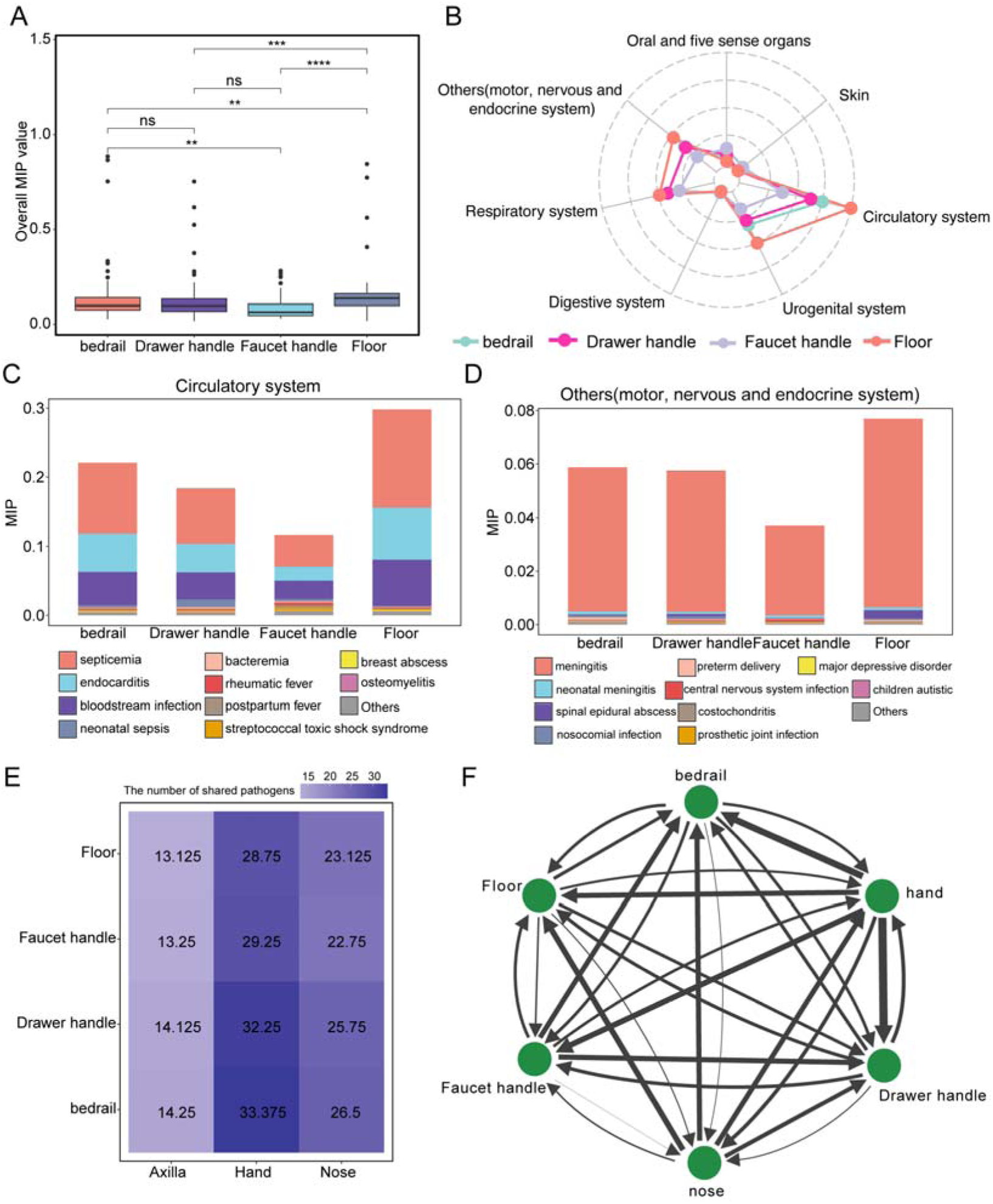
Pathogen detection and risk assessment in hospital wards using MIP. (A, B) The overall MIP value (A) and the potential risk pattern of human diseases (B) in four types of hospital environmental samples. (C, D) The diseases related to circulatory system (C) and others (D) in hospital wards. (E) Shared pathogen heatmap for patients and hospital wards. (F) The potential transmission route of pathogen in hospital wards using microbial source tracking, the contribution values from source sites to query sites were represented by the width of the edges.

Additionally, to understand the changes in potential pathogenic risks posed by microorganisms in different locations of the hospital wards following its opening, we compared the MIP values between preopening and post-opening periods. The results indicated that the overall MIP values of bedrails in post-opening were significantly higher than preopening (Supplementary Fig. 4A). However, the overall MIP values of drawer handles, floors, and faucet handles between preopening and post-opening periods did not have significantly differences (Supplementary Fig. 4B, C, D). These findings indicated that, despite floors possessed the highest potential pathogenic risk among all sites, its risk level remained relatively stable before and after the hospital opening. Conversely, bedrails exhibited a significantly increase in potential pathogenic risk after hospital opening. After the hospital opening, pathogens such as *Prevotella melaninogenica*, *Streptococcus cristatus*, and *Enterococcus faecalis* in bedrails were significantly enriched than preopening, which result in the potential disease risks related to the OFSO and SK in bedrails significantly increased after hospital opening (Supplementary Fig. 4E-K). After hospital opening, the potential pathogenic risk related to OFSO, such as streptococcal sore throat (Supplementary Fig. 4L), the potential pathogenic risk related to SK, such as botryomycosis, necrotizing fasciitis, cellulitis, and erythrasma, were both showed a significantly increase (Supplementary Fig. 4M).

To further investigate potential pathogen transmission pathways, we analyzed the overlap of pathogens identified both in patients and hospital wards. As shown in Fig. 5E, the most frequent communication between hospital wards and patient body was patient hands to bedrail. Furthermore, by mapping the pathogenic transmission network within hospital wards, we found that patient hands were major hub for pathogenic transmission (Fig. 5F). Pathogens were primarily transmitted from patient hands to various other sites, including bedrails, drawer handles, floors, and faucet handles. This transmission pattern is attributed to several factors, such as the frequent direct contact between patient hands with contaminated surfaces, the ability of pathogens to persist and survive on human skin, and the potential for secondary transfer of pathogens to other surfaces and individuals. In summary, these results suggested that patients were the primary reservoirs of the environmental pathogens within hospital wards, in the future, hand hygiene and environmental cleaning in reducing the spread of infections in healthcare settings is critical importance.

## Discussion

In this study, we used high-quality 16S rRNA gene amplicon sequences to explore bacterial colonization and transmission dynamics in a newly opened hospital.

Bacterial communities displayed clear spatial patterns, with significant shifts in both alpha and beta diversity after hospital opening. Patient microbiomes, particularly from hands, strongly influenced the composition, structure, and abundance of bacterial communities in hospital wards, highlighting the critical role of human activity in structuring hospital microbiomes. Additionally, we mapped the microbial transmission network and evaluated the change of the potential disease risks across key sites within hospital wards after opening. These findings will provide a scientific basis for the development of more effective strategies to resolve hospital-acquired infections.

Understanding how microbial communities colonize and persist on built-environment surfaces is essential for assessing potential public health risks^2,46^. A recent study investigated microbial colonization in the preterm infant gut and found that strains detected in hospitalized infants also occur in sinks and on surrounding surfaces, including *Staphylococcus epidermidis*, *Enterococcus faecalis*, *Pseudomonas aeruginosa*, and *Klebsiella pneumoniae*^47,48^, therefore, hospital wards are reservoirs for early microbiome colonizers. Kang et al. used metagenomic analysis to investigate the microbiome of skin and metro system, they found microbial communities in metro system are decidedly shaped by human interactions and traffic flows^8^. Additionally, Evidence from longitudinal studies demonstrates that dust microbiomes undergo significant temporal changes. In kindergartens, human-associated bacteria not only vary in abundance but also include resistant strains, raising concerns about the spread of resistance^6^. Another longitudinal study of the indoor microbiome found that the indoor microbiome of the new-dwellings was rapidly colonized by its occupants^46^.

Similar dynamics has been observed in hospitals, where microbial exchange occurs between patients and their surrounding surfaces.^11,26^. Our longitudinal study on the microbial colonization of hospital surfaces also showed consistent shifts in bacterial composition and diversity across all tested sites. Specifically, in our study, we found Proteobacteria, Firmicutes, Actinobacteria, and Bacteroidetes were dominant in hospital wards (Fig.1A), these four phyla were also found to be dominant in other building environment, such as airplane cabins^49^, hotel rooms^50^, kindergartens^6^, and dormitories^51^. Additionally, the total relative abundance of these four phyla on human skin was higher than 99%^52^. At the genus level, we identified *Acinetobacter* as the dominant genus on floors and showing a significant decline following patient occupancy (Fig.2A, B). Human-associated and potentially clinically significant bacteria (e.g., *Staphylococcus*, *Corynebacterium*, and *Streptococcus*) significantly increased in relative abundances after patient occupancy (Supplementary Fig. 1A, B). These results indicated that patient occupancy influences the composition of bacterial community in hospital wards.

In addition, we found that the alpha diversity of floors and drawer handles in preopening was significantly lower than after-opening, in contrast, the alpha diversity of bedrails and faucet handles was significantly higher in preopening than after-opening (Fig. 2C, D). Previous studies also showed a higher diversity of microbial communities in floor, doorhandle and sink samples after opening^11^, however, another study indicated the alpha diversity of floor samples did not significantly change between preopening and after-opening^26^. These differences may be partially explained by the complex relationship between microbial communities dynamic and human action, and cleaning and disinfection. The floors and drawer handles are exposed to more human activities after opening compared to before opening. With frequent personnel movement and the operation of medical equipment, these surfaces are more likely to come into contact with diverse microbial sources, leading to an increase in the alpha diversity of bacterial communities. The reduced alpha diversity observed on bedrails and faucet handles may be attributed to the stricter cleaning and disinfection protocols implemented in these areas. These surfaces are frequently touched by both patients and healthcare workers, making them focal points for targeted cleaning and disinfection to minimize the risk of pathogen transmission. Frequent cleaning and disinfection can eliminate certain microbes, thereby reducing microbial diversity in these areas.

Although routine disinfection and strict isolation measures in hospital wards reduce bacterial dissemination, the frequent occurrence of co-infections highlights the possibility of bacterial transmission within the hospital environment^53^. Using the SourceTracker tool^41^, our study elucidated bacterial transmission patterns in hospital wards and found that bacteria located in patients and sites of hospital wards could easily interact. Patient hands were the major hub for bacterial transmission in hospital wards (Fig.4A). Bacteria from patient hands spread to sites in hospital wards that are directly touched by patients, such as bedrails, drawer handles, faucet handles. A study investigated the bacterial transmission using 124 medical staff in different hospital departments, and found mobile phones are a potential transmission vector^54^. In addition, researcher found that bedrails and inside floor in hospital are important site of bacterial transmission^27^. Furthermore, some bacteria with high transfer capacity pose a greater risk of infecting humans. Therefore, transfer capacity of microbes also deserved attention. In our study, we found that some non-pathogenic genera were widely present in hospital wards, such as *Lactobacillus*, and *Cornebacterium*. Further research on the impact of these genera in the hospital setting is warranted. Additionally, we identified several pathogens with great transfer capacity, such as *Staphylococcus* and *Streptococcus*. These two genera not only had a high transfer capacity, but also exhibit high relative abundance in hospital wards. These results highlight the potential risk of co-infection in hospital wards.

By employing network analyses, we characterized the co-occurrence pattern of Transfer Easy taxa in hospital wards, and found that the interactions between Transfer Easy taxa were frequent, and only a minority of Transfer Easy taxa had connections to other nodes in hospital wards (Fig.4C). The tight internal interaction patterns observed among Transfer Easy taxa may give them population resistance and transmission advantages^55,56^. This structural feature reflects the selection pressures imposed by hospital environment and highlights their unique ecological strategy. Transfer Easy taxa gain advantages in dynamic and competitive environments by reducing direct competition with other microbes, instead optimizing their capacity for transmission and colonization. This phenomenon not only underscores the complexity and adaptability of microbial communities in clinical settings, but also provides a new perspective for preventing hospital-acquired infections. Future research should combine multiple omics data and experimental validation to deeply elucidate their interaction mechanisms and clinical significance.

## Conclusions

In conclusion, this study investigates the colonization and transmission dynamics of the bacterial community in hospital wards, revealing the significant influence of patient occupancy on the bacterial community in hospital wards. Furthermore, our study explores bacterial transfer capacity and potential disease risk of different sites in hospital wards, and provides suggestion for the key sites that require enhanced disinfection and protection in the future. These findings provide valuable insights to guide the prevention and control of co-infections in hospital settings.

## Acknowledgments

We thank the members at Beijing Tiantan Hospital, Capital Medical University for the critical discussions of this work. In addition, we thank the High-performance Computing Platform of Peking University for providing the computing platform.

## Author Contributions

All authors contributed intellectual input and assistance to this study. L.Y.A. and J.S.C. conceived the study and performed the original analysis. X.N.L., X.L., and Y.X.C. collected raw data. L.Y.A. and Y.N. co-wrote the paper. Q.L., X.Q.S., and J.S.C revised it. Y.N. and L.Y.A. raised the funding for the project. All authors discussed the results and commented on the article.

## Ethics approval

Not applicable.

## Consent to participate

Not applicable.

## Consent for publication

Not applicable.

