## Supplementary information for "Longitudinal analysis of bacterial community dynamics in normal hospital ward following patient occupancy"

\# Corresponding authors:

**Supplemental figures**
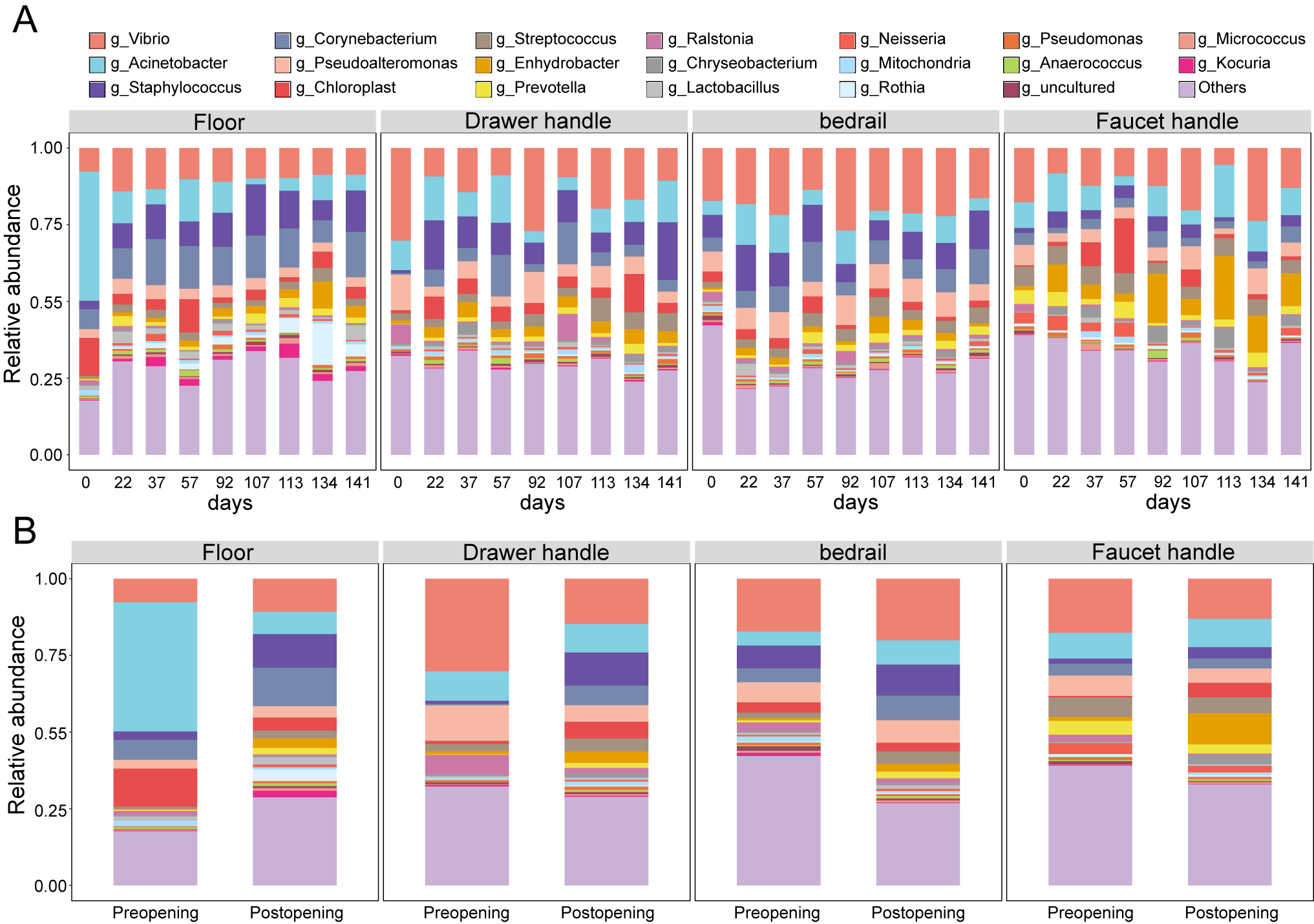

Supplementary Fig. 1 Changes in the relative abundance of top10 genera with time (A), and before and after hospital opening (B).

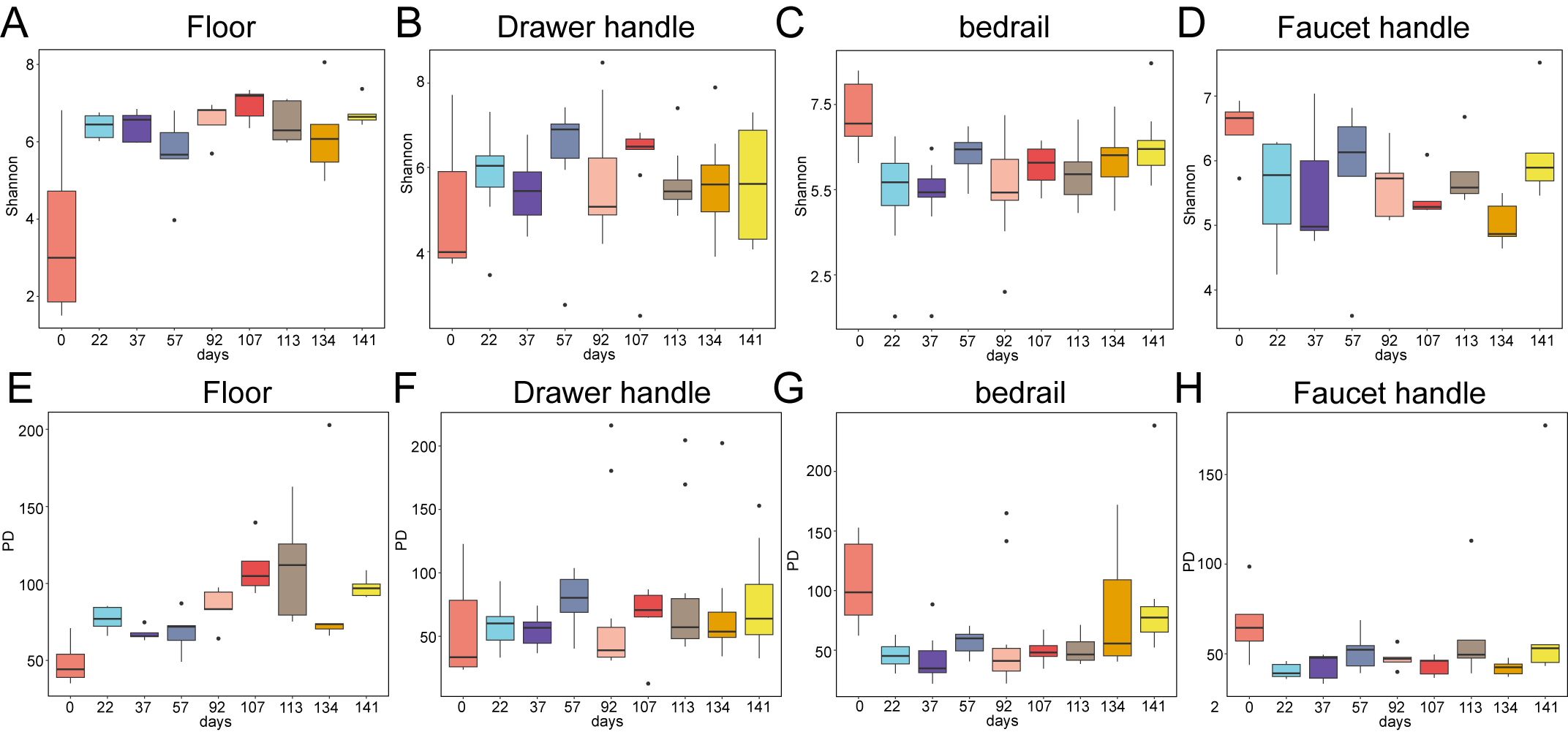

Supplementary Fig. 2 The changes in the Shannon diversity (A, B, C, D) and Faith’s PD (E, F, G, H) index of samples after hospital opening.

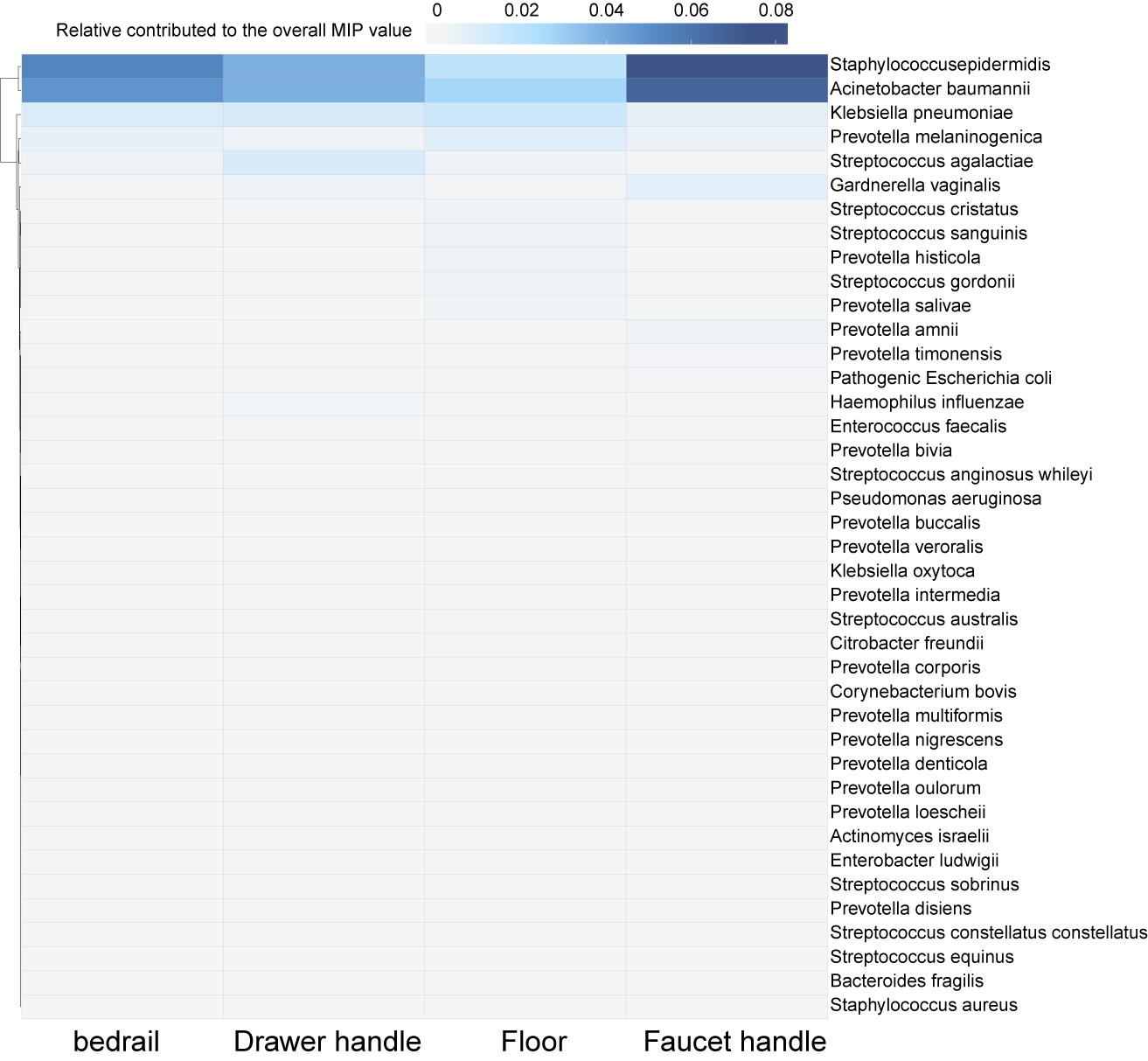

Supplementary Fig. 3 Heatmap showed the relative abundance (> 0.01%) of pathogens in hospital ward. The color indicated relative abundance.

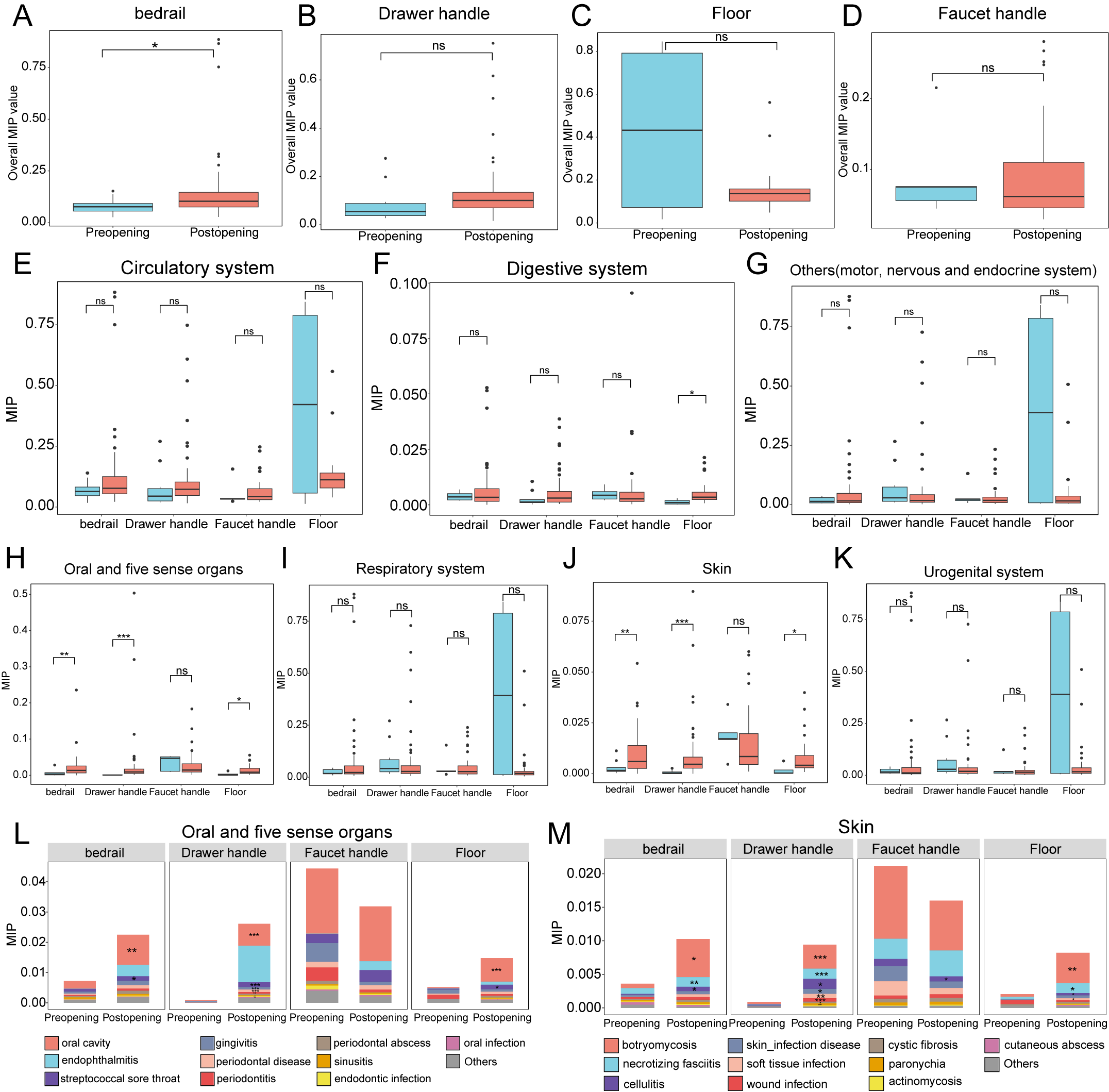

Supplementary Fig. 4 The change of potential pathogenic risks after hospital opening. (A, B, C, D) The overall MIP value in bedrails (A), drawer handles (B), floor (C), and faucet handles (D). (E, F, G, H, I, J, K) The changes in potential pathogenic risks associated with the circulatory system (E), digestive system (F), others (e.g., motor system, nervous system, and endocrine system) (G), oral and five sensory organs (H), respiratory system (I), skin (J), and urogenital system (K) after hospital opening. (L, M) The change of diseases related to oral and five sensory organs (L), and skin (M) after hospital opening.
